# Single-molecule observation of ligand binding and conformational changes in FeuA

**DOI:** 10.1101/624817

**Authors:** Marijn de Boer, Giorgos Gouridis, Yusran Abdillah Muthahari, Thorben Cordes

## Abstract

The specific binding of ligands by proteins and the coupling of this process to conformational changes are fundamental to protein function. We designed a fluorescence-based single-molecule assay and data analysis procedure that allows the simultaneous real-time observation of ligand binding and conformational changes in FeuA. The substrate-binding protein FeuA binds the ligand ferri-bacillibactin and delivers it to the ABC importer FeuBC, which is involved in iron uptake in bacteria. The conformational dynamics of FeuA was assessed via Förster resonance energy transfer (FRET), whereas the presence of the ligand was probed by fluorophore quenching. We reveal that ligand binding shifts the conformational equilibrium of FeuA from an open to a closed conformation. Ligand binding occurs via an induced-fit mechanism, i.e., the ligand binds to the open state and subsequently triggers a rapid closing of the protein. However, FeuA also rarely samples the closed conformation without the involvement of the ligand. This shows that ligand interactions are not required for conformational changes in FeuA. However, ligand interactions accelerate the conformational change 10000-fold and temporally stabilize the formed conformation 250-fold.

**SIGNIFICANCE STATEMENT:** Ligand binding and the coupling of this process to conformational changes in proteins are fundamental to their function. We developed a single-molecule approach that allows the simultaneous observation of ligand binding and conformational changes in the substrate-binding protein FeuA. This allows to directly observe the ligand binding process, ligand-driven conformational changes as well as rare short-lived conformational transitions that are uncoupled from the ligand. These findings provide insight into the fundamental relation between ligand-protein interactions and conformational changes. Our findings are, however, not only of interest to understand protein function, but the developed data analysis procedure allows the determination of (relative) distance changes in single-molecule FRET experiments, for situations in which donor and acceptor fluorophore are influenced by quenching processes.

## INTRODUCTION

The non-covalent and specific interactions between ligands and proteins underlies almost all biological processes. The coupling of these binding events to conformational changes allows proteins to act as highly efficient enzymes, signal transducers, motors, switches or pumps^1^. Two basic models that describe the coupling between protein conformational changes and ligand binding are the induced-fit^2^ and conformational selection mechanism^3^. These mechanisms represent the two limiting pathways on the energy landscape that connect unliganded and liganded conformational states (Fig. 1). In the induced-fit mechanism, ligand interactions trigger conformational changes, whereas in the conformational selection mechanism, ligand interactions selectively stabilize a subset of conformations that are already present in the unliganded protein (Fig. 1). Both mechanisms require intermediate states that are formed during the ligand-binding process. For example, when a protein switches between two conformational states, such as an open and closed conformation (Fig. 1), an open-liganded state in the induced-fit mechanism or a closed-unliganded state in the conformational selection mechanism, are essential intermediate states. However, the study of such transient and thermodynamically unstable states remains experimentally challenging.

**Figure 1.**
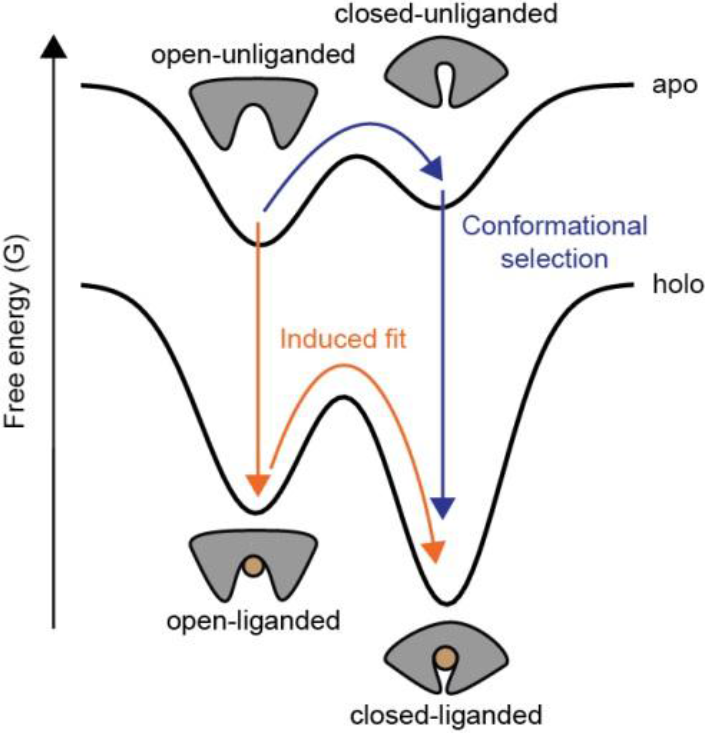
Conformational landscape and ligand-binding mechanism of FeuA. Energy landscape of a protein adopting open and closed conformations in the absence (apo) and presence of ligand (holo). Ligand recognition can be coupled to conformational changes by an induced-fit mechanism, wherein substrate binding facilitates a conformational change, or a conformational selection mechanism, in which thermal energy induces conformational changes to enable ligand recognition.

Primarily driven by high resolution structural analysis of proteins adopting different conformations in the absence and presence of ligand, an induced-fit mechanism was implied for many proteins. However, advancement in especially single-molecule spectroscopy^4–7^, NMR^8–10^ and other spectroscopic methods^11^, revealed that the structure of proteins are highly flexible and can undergo large conformational changes intrinsically, i.e., in the absence of the ligand molecule. Examples are dihydrofolate reductase^12^, adenylate kinase^9^, ubiquitin^13^, ABC exporters^7^, substrate-binding proteins of Type I ABC importers^4^, DNA polymerase^14, 15^ and RNase A^10^. Due to the occurrence of these intrinsic conformational changes, a conformational selection mechanism has been proposed to underlie the ligand binding process of many protein systems^3^. However, the unambiguous determination of the binding mechanism requires simultaneous monitoring of the ligand binding events and the protein conformation. Moreover, the intrinsic conformational ensemble can in principle be investigated by studying the protein in the absence of ligand. However, trace contaminations of ligand can make this assessment experimentally difficult, especially under single-molecule conditions. To bypass these problems, we here established a fluorescence-based single-molecule assay and data analysis procedure that allows the simultaneous real-time observation of conformational changes and ligand binding in FeuA.

The substrate-binding protein FeuA is associated with the ATP-binding cassette (ABC) transporter FeuBC from *Bacillus subtilis*^16^. This type II ABC importer is involved in the uptake of Fe^3+^ ions by ATP-driven active transport of the siderophore bacillibactin (BB) in complex with Fe^3+^ (FeBB)^16^. FeuA and other structurally related substrate-binding proteins (SBPs^17^) or domains (SBDs^17^) represent the primary receptors of bacterial ABC importers^17^, tripartite ATP‐independent periplasmic (TRAP) transporters^18^, and others^19^. These proteins capture the ligand from the external environment for delivery to the membrane transporter complex and import into the cell. The structure of FeuA has a characteristic SBP-fold^19^, consisting of two subdomains connected by a linking region, with ligand binding occurring at the interface of the subdomains (Fig. 2a)^20^. Crystallography studies suggest that FeuA undergoes conformational changes that involves a domain reorientation that engulfs the ligand, leading to opening and closing of the protein^20^. This apparently simple binary conformational switch, which is involved in molecular recognition, is investigated in this paper to obtain insight into the relation between ligand-interactions and the coupling to protein conformational changes.

**Figure 2.**
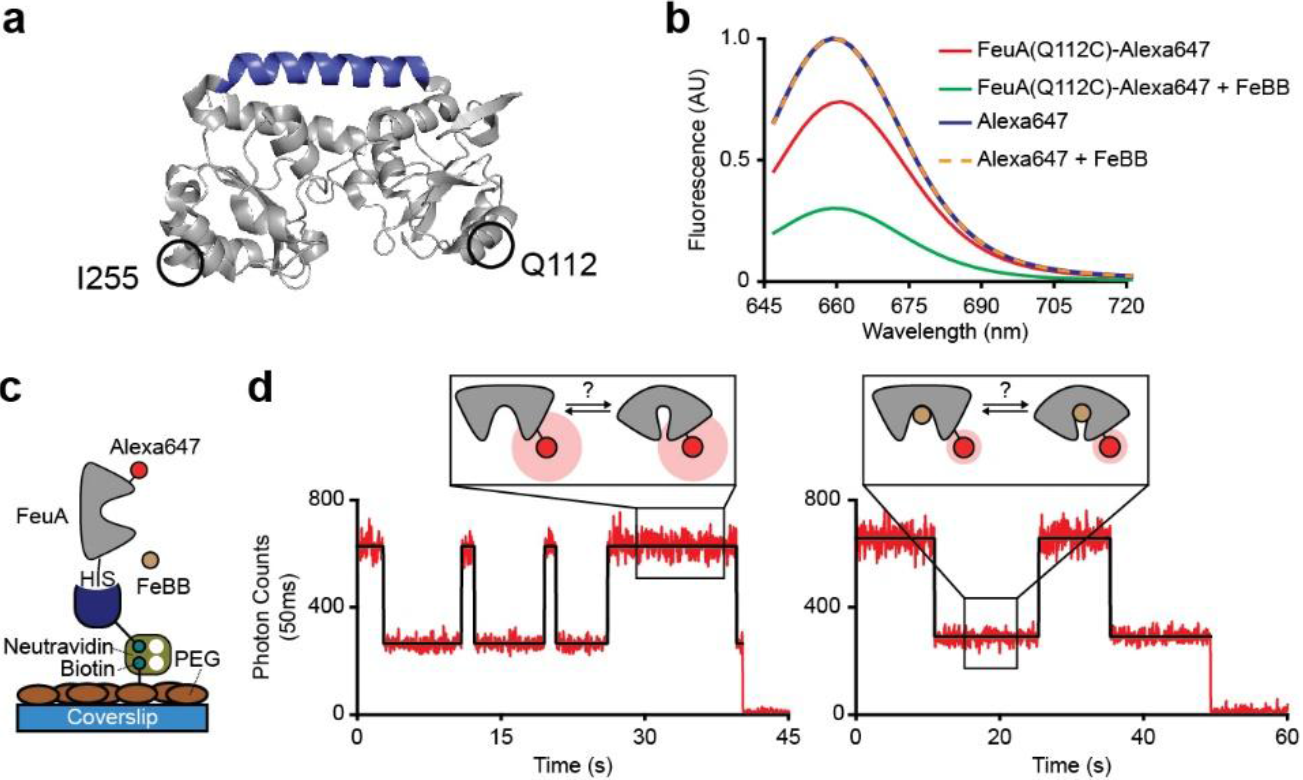
Direct observation of ligand binding in FeuA using fluorescence quenching. **a**, X-ray crystal structure of the open-unliganded state of FeuA (PBD ID: 2WI8). Hinge region is indicated in blue. **b**, Emission spectra of FeuA labeled with Alexa647 and free Alexa647 in the presence and absence of 5 μM FeBB. **c**, Schematic of the experimental strategy to study individual surface-tethered FeuA proteins. **d**, Representative fluorescence trajectories of FeuA(Q112C)-Alexa647 in the presence of 40 nM FeBB. The fluorescent intensity (red) with the most probable state-trajectory of the Poisson Hidden Markov Model (PHMM) (black) are shown. The number of analysed molecules is provided in Table S3.

## MATERIALS AND METHODS

### Gene isolation, protein expression and purification

The *feu*A gene (Uniprot: P40409) was isolated by PCR from the genome of *Bacillus subtilis* subsp. *subtilis* str. *168*. The primers were designed to exclude the signal peptide (amino acids 1-19), and cysteine 20 (which is probably post-translationally lipidated) with *Nde*I/*Hind*III restriction sites. Primers are indicated in Table S1. The generated PCR fragment was A-tailed and ligated into the PGEM-T Easy Vector System (Promega)^21^. After removing the *Nde*I restriction site internal to the *feu*A gene by a silent mutation, the gene was sub-cloned in the pET20b vector (Merck) using the *Nde*I/*Hind*III sites. Protein derivatives including the cysteine and the silent mutation were constructed using QuickChange mutagenesis^22^. All sequences were checked for correctness by sequencing.

Cells harbouring plasmids expressing the FeuAHis_6_ wild-type and derivatives were grown at 37ºC until an optical density (OD_600_) of 0.5 was reached. Protein expression was then induced by addition of 0.25 mM isopropyl β-D-1-thiogalactopyranoside (IPTG). After 3 hours of induction cells were harvested. DNase 500 ug/ml (Merck) was added and passed twice through a French pressure cell at 1,500 psi. 2 mM phenylmethylsulfonyl fluoride (PMSF) was added to inhibit proteases. The soluble supernatant was isolated by centrifugation at 50,000×*g* for 30 min at 4 °C. The soluble material was purified and loaded on Ni^2+-^sepharose resin (GE Healthcare) in 50 mM Tris-HCl, pH 8.0, 1 M KCl, 10% glycerol, 10 mM imidazole, 1 mM dithiothreitol (DTT). The immobilized proteins were washed (50 mM Tris-HCl, pH 8.0, 50 mM KCl, 10 % glycerol, 10 mM imidazole, 1 mM DTT and subsequently with 50 mM Tris-HCl, pH 8.0, 1 M KCl, 10 % glycerol, 30 mM imidazole, 1 mM DTT) and eluted (50 mM Tris-HCl, pH 8.0, 50 mM KCl, 10 % glycerol, 300 mM imidazole, 1 mM DTT). Protein fractions were pooled (supplemented with 5 mM EDTA, 10 mM DTT), concentrated (10.000 MWCO Amicon; Merck-Millipore), dialyzed against 100-1000 volumes of buffer (50 mM Tris-HCl, pH 8.0, 50 mM KCl, 50% glycerol, 10 mM DTT), aliquoted and stored at −20°C until required.

### Protein labelling

Labelling was performed with the maleimide dyes Alexa555 and Alexa647 (ThermoFisher). The purified proteins were treated with 10 mM DTT for 30 min at 4°C to reduce oxidized cysteines. The protein sample was diluted to 1 mM DTT, immobilized on a Ni^2+^-Sepharose resin (GE Healthcare) and washed with 10 column volumes of buffer A (50 mM Tris-HCl, pH 7.4, 50 mM KCl). The resin was incubated 2-8 hrs at 4°C with the dyes dissolved in buffer A. The molar dye concentration was 20-times higher than the protein concertation. Unbound dyes were removed by washing the column with 20 column volumes of buffer A and eluted with 400 mM imidazole. The labelled proteins were further purified by size-exclusion chromatography (Superdex 200, GE Healthcare) using buffer A. The sample composition was assessed by absorbance measurement at 280 nm (protein), 559 nm (Alexa555), and 645 nm (Alexa647) to determine labelling efficiency. For all samples the labelling efficiency was >90%.

### Ensemble fluorescence measurements

The fluorescence spectra were recorded on a scanning spectrofluorometer (Jasco FP-8300). Emission spectra were recorded by excitation at 635 nm (5 nm bandwidth) in steps of 2 nm (2 nm emission bandwidth and 8 s integration time). Fluorescence anisotropy values *r* = (*I*_*VV*_ − *GI*_*VH*_)/(*I*_*VV*_ + 2*GI*_*VH*_) were determined around the emission maxima of the two fluorophores (for donor, λ_ex_ = 535 nm and λ_em_ = 580 nm; for acceptor, λ_ex_ = 635 nm and λ_em_ = 660 nm; 5 nm bandwidth and 8 s integration time). *I*_*VV*_ and *I*_*VH*_ are the fluorescence emission intensities in the vertical and horizontal orientation, respectively, upon excitation along the vertical orientation. The correction factor is *G* = *I*_*HV*_/*I*_*HH*_, where *I*_*HV*_ and *I*_*HH*_ are the fluorescence emission intensities in the vertical and horizontal orientation, respectively, upon excitation along the horizontal orientation. Experiments were performed in buffer A at a concentration of 100-250 nM of labelled proteins and free-fluorophores at room temperature.

### Solution-based smFRET and ALEX

Solution-based smFRET and alternating laser excitation (ALEX)^23, 24^ experiments were carried out at 25-100 pM of labeled protein at room temperature in buffer A in the absence or presence of ferri-bacillibactin (EMC biochemicals). Microscope cover slides (no. 1.5H precision cover slides, VWR Marienfeld) were coated with 1 mg/mL BSA (Merck) for 1 min and unbound BSA was subsequently removed by washing with buffer A.

The measurements were performed using a home-built confocal microscope as described before^4^. In brief, two laser-diodes (Coherent Obis) with emission wavelength of 532 and 637 nm were modulated in alternating periods of 50 μs. The laser beams where coupled into a single-mode fiber (PM-S405-XP, Thorlabs) and collimated (MB06, Q-Optics/Linos) before entering a water immersion objective (60X, NA 1.2, UPlanSAPO 60XO, Olympus). The fluorescence was collected by excitation at a depth of 20 μm. Average laser powers where 30 μW at 532 nm (~30 kW/cm^2^) and 15 μW at 637 nm (~15 kW/cm^2^). Light was separated by a dichroic beam splitter (zt532/642rpc, AHF Analysentechnik), mounted in an inverse microscope body (IX71, Olympus). Emitted light was focused onto a 50 μm pinhole and spectrally separated (640DCXR, AHF Analysentechnik) onto two single-photon avalanche diodes (TAU-SPADs-100, Picoquant) with appropriate spectral filtering (donor channel: HC582/75; acceptor channel: Edge Basic 647LP; AHF Analysentechnik). Registration of photon arrival times and alternation of the lasers was controlled by an NI-Card (PXI-6602, National Instruments).

### Analysis of solution-based smFRET data

Photons were binned in 1 ms intervals and only bins with a total of >200 photons considering all detection channels were further analyzed. Three photon count rates were measured: 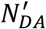 (acceptor emission upon donor excitation), 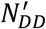 (donor emission upon donor excitation) and 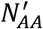 (acceptor emission upon acceptor excitation)^23^. The background counts were estimated by excluding all time-bins containing more than 20 counts and calculating the mean count rate over all remaining time-bins. The leakage and direct excitation contributions were determined from the donor- and acceptor-only labeled molecules as described by Lee et al^25^. Crosstalk and background correcting 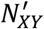 yields *N*_*XY*_. The proximity FRET efficiency *E*_*PR*_ is

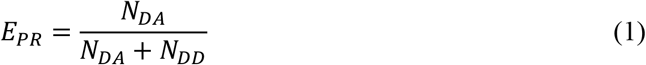

and the Stoichiometry *S* is

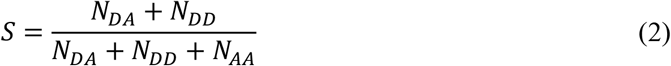

The appropriate sub-populations within *E*_*PR*_ and *S* dataset where clustered using a Gaussian Mixture Model, with one (apo) or two (holo) multivariate normal distributions. Molecules were assigned to the component yielding the highest posterior probability and are within 98% of the probability mass. For each cluster the average *N*_*DA*_/*N*_*DD*_, *N*_*DD*_/*N*_*DA*_ and *E*_*PR*_ was calculated. All post-processing steps were programmed in Matlab (MathWorks).

### Theory of interprobe distance ratio estimation

The distance between the donor and acceptor fluorophore *r* is related to the FRET efficiency *E* via

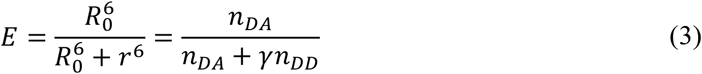

where *R*_0_ is the Föster radius, *n*_*DD*_ and *n*_*DA*_ are the background- and spectral crosstalk-corrected donor and acceptor emission count rates when the donor is excited, respectively. *γ* = *ϕ_A_η_D_*/*ϕ_D_*η_A_** depends on the donor and acceptor quantum yields, *ϕ_D_* and *ϕ_A_*, respectively, and the detection efficiencies of the donor- and acceptor emission detection channels, *η_D_* and *η_A_*, respectively^25^. Equation (3) can be rewritten as

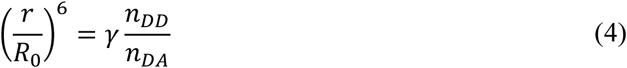

Let *r*_1_ and *r*_2_ denote the (average) donor and acceptor fluorophore distance of two states. By using the definition of *γ* and noting that 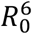 is proportional to *ϕ_D_*, we find that the ratio between *r*_1_ and *r*_2_ satisfies:

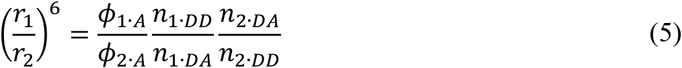

where *n*_*i·DA*_ and *n*_*i·DD*_ are the donor and acceptor count rates when the donor is excited and belong to *r*_*i*_, and *ϕ*_*i·A*_ is the acceptor quantum yield. Let us now consider how the distance ratio (*r*_1_/*r*_2_)^6^ can be estimated from the data. We use the following notation: *N*_*i·XY*_ represents the measured count rate of *Y* emission upon *X* excitation belonging to *r*_*i*_. *N*_*i·XY*_ is corrected for spectral crosstalk and background. We can assume that the relaxation times of the excited states of the fluorophores are short compared to the time between two consecutively detected photons, so that there is no correlation between consecutive photons and the distribution of *N*_*i·XY*_ can be approximated by a Poisson distribution^26^. Then,

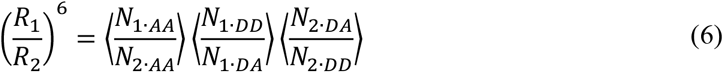

with

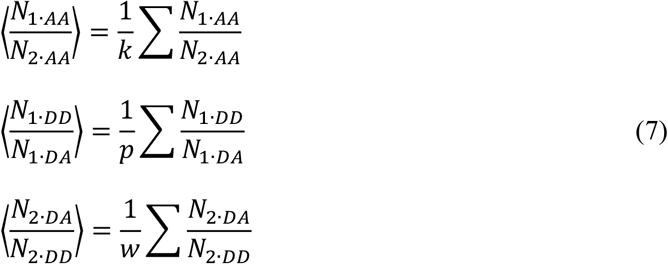

where *k*, *p* and *w* denote the number of observations, is an unbiased and consistent estimator for (*r*_1_/*r*_2_)^6^. The sum in equation (7) extends over all observations, i.e. the total number of traces or time-bins (see results section for details). In the absence of fluorophore quenching by the ligand we find that *ϕ*_*F·A*_ = *ϕ*_*B·A*_ and

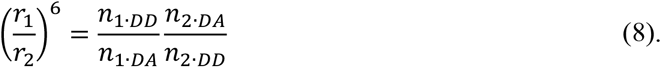

So, an unbiased and consistent estimator for equation (8) is

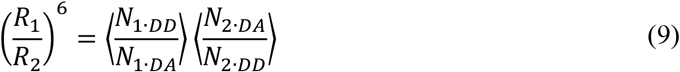

Further details and a full derivation are provided in the Supplementary Information.

### Scanning confocal microscopy

Confocal scanning microscopy was performed at room temperature using a home-built confocal scanning microscope as described before^27, 28^. In brief, surface scanning was performed using a XYZ-piezo stage with 100×100×20 μm range (P-517-3CD with E-725.3CDA, Physik Instrumente). The detector signal was registered using a HydraHarp 400 picosecond event timer and a module for time-correlated single photon counting (both Picoquant). Data were recorded with constant 532 nm excitation at intensities between 0.1 and 5 μW (~25-1250 W/cm^2^) for smFRET measurements and 0.1 μW (~25 W/cm^2^) when the protein was only labeled with Alex647 (640 nm excitation) or Alex555 (532 nm excitation). Scanning images of 10×10 μm were recorded with 50 nm step size and 1 ms integration time at each pixel. After each surface scan, the positions of labeled proteins were identified manually. Surface immobilization was done and a flow-cell arrangement was prepared as described before^28, 29^. Measurements were done in buffer A supplemented with 1 mM 6-hydroxy-2,5,7,8-tetramethylchroman-2-carboxylic acid (Trolox; Merck) and 10 mM Cysteamine (Merck).

### Analysis of fluorescence trajectories

Fluorescence trajectories were recorded in time-bins of varying length as stated in the text or figure captions. We use the following notation: 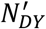 is the uncorrected count rate, 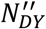 is the background corrected count rate and *N*_*DY*_ the background and spectral crosstalk corrected count rate of *Y* emission (*<u>D</u>*onor, *<u>A</u>*cceptor) upon donor excitation. The apparent FRET efficiency is 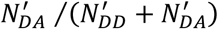. Only traces lasting longer than 20 time-bins, having on average more than 10 photons per time-bin which showed clear bleaching steps were used for further analysis.

Equation (9) was used to estimate the interprobe distance ratio, with 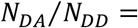 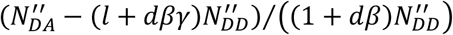 and 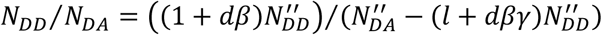 where *l*, *d*, *γ* and *β* are correction factors^25^. Background was determined as the average count rate per channel when the fluorophores have bleached. The correction factors were determined using solution-based ALEX as described in Lee et al^25^. In brief, the *l* and *d* factors were determined from the donor-and acceptor-only labeled FeuA molecules and the *β* and *γ* factors using a protein reference standard (MalE^4^). All correction factors were determined on the same microscope also used for the surface-based measurements (details of setup in Gouridis et al.^28^).

The state-trajectories were modelled by a Hidden Markov Model (HMM)^30^. For this an implementation of HMM was programmed in Matlab (MathWorks) as described previously^4^. We assume that the FRET and acceptor-only and donor-only fluorescence trajectory can be considered as a HMM with only two states having a one-dimensional Gaussian- or a Poisson-output distribution, respectively. The Gaussian distribution of state *i* (*i* =1, 2) is defined by the average and variance. The Poisson distribution of state *i* (*i* =1, 2) is defined by the average intensity of the acceptor or donor in state *i*. The likelihood function was maximized by using the Baum-Welch algorithm^31^. The most probable state-trajectory was found using the Viterbi algorithm^32^. The time spend in each state was interfered from the most probable state-trajectory and used to construct a 95% confidence interval for the mean lifetime. The quenching ratio for each molecule was obtained by taking the ratio of the intensity levels as obtained from the Poisson HMM.

## RESULTS

### Direct observation of binding and unbinding of ligand

To investigate ligand binding of FeuA at the single-molecule level we labelled FeuA with the fluorophore Alexa647 in one of its subdomains, by introducing a single cysteine residue at a non-conserved position, which is solvent-exposed and distant from the binding pocket (Q112C; Fig. 2a). First, we determined the emission spectra of FeuA-Alexa647 and free Alexa647 in the presence and absence of FeBB. We observed that only the fluorescence intensity of FeuA-Alexa647 was quenched in the presence of 5 μM FeBB (Fig. 2b). Since no quenching was observed for free Alexa647 (Fig. 2b), we attribute this to binding of FeBB by FeuA.

To directly observe ligand binding and unbinding events, the fluorophore emission of individual surface-tethered labeled proteins was studied over time by using confocal scanning microscopy (Fig. 2c). Representative fluorescent intensity trajectories of FeuA in the presence of 40 nM FeBB are shown in Fig. 2d. All analyzed fluorescence trajectories show a single bleaching step, indicating that single molecules are examined (Fig. 2d). Only in the presence of FeBB we observed stochastic switching between two intensity levels (Fig. 2d), caused by fluorescence quenching of Alexa647 by FeBB. Thus the intensity fluctuations can be interpreted as individual binding and unbinding events of the ligand FeBB to FeuA. To substantiate this claim, we determined the relative population of the lower intensity level to estimate the dissociation constant K_D_ of FeBB binding by FeuA. From the analysis of 50 traces we obtained an estimated K_D_ of 20 nM, which is in good agreement with the value obtained from tryptophan fluorescence (K_D_ = 27 ± 1 nM)^33^. In summary, our assay can directly probe the ligand binding process of FeuA. However, how the binding and unbinding events are coupled to the conformational changes in FeuA remains unclear.

### Conformational states of FeuA

To further elucidate this, we used Förster resonance energy transfer (FRET) to analyse the conformational changes of FeuA. In the assay, each of the two subdomains was stochastically labelled with either a donor (Alexa555) or an acceptor fluorophore (Alexa647). Surface-exposed and non-conserved residues, showing largest distance changes according to the crystal structures of the open and closed states^20^, were chosen as cysteine positions for fluorophore labelling (Q112C/I255C; Fig. 2a). The relationship between FRET efficiency and interprobe distance requires free fluorophore rotation, which was verified by steady-state anisotropy measurements (Table S2).

We used confocal microscopy with alternating laser excitation (ALEX)^23^ to explore the conformational states of individual freely diffusing proteins (Fig. 3a). During its diffusional transit through the excitation volume of a confocal microscope, the labelled protein generates short fluorescent bursts, allowing the determination of the apparent FRET efficiency and the stoichiometry *S* (see Materials and Methods section for details). To retrieve interprobe distances, the apparent FRET efficiency was corrected for background and spectral crosstalk to obtain the proximity ratio *E*_*PR*_. In our assays, changes in the apparent FRET efficiency and *E*_*PR*_ can originate from interprobe distance changes, but also due to quenching of the fluorophores by binding of FeBB (Fig. 3b). Finally, *S* relates the total fluorescence recorded after donor excitation in the green and red detection channel to the total fluorescence after direct donor and acceptor excitation in each detection channel.

**Figure 3.**
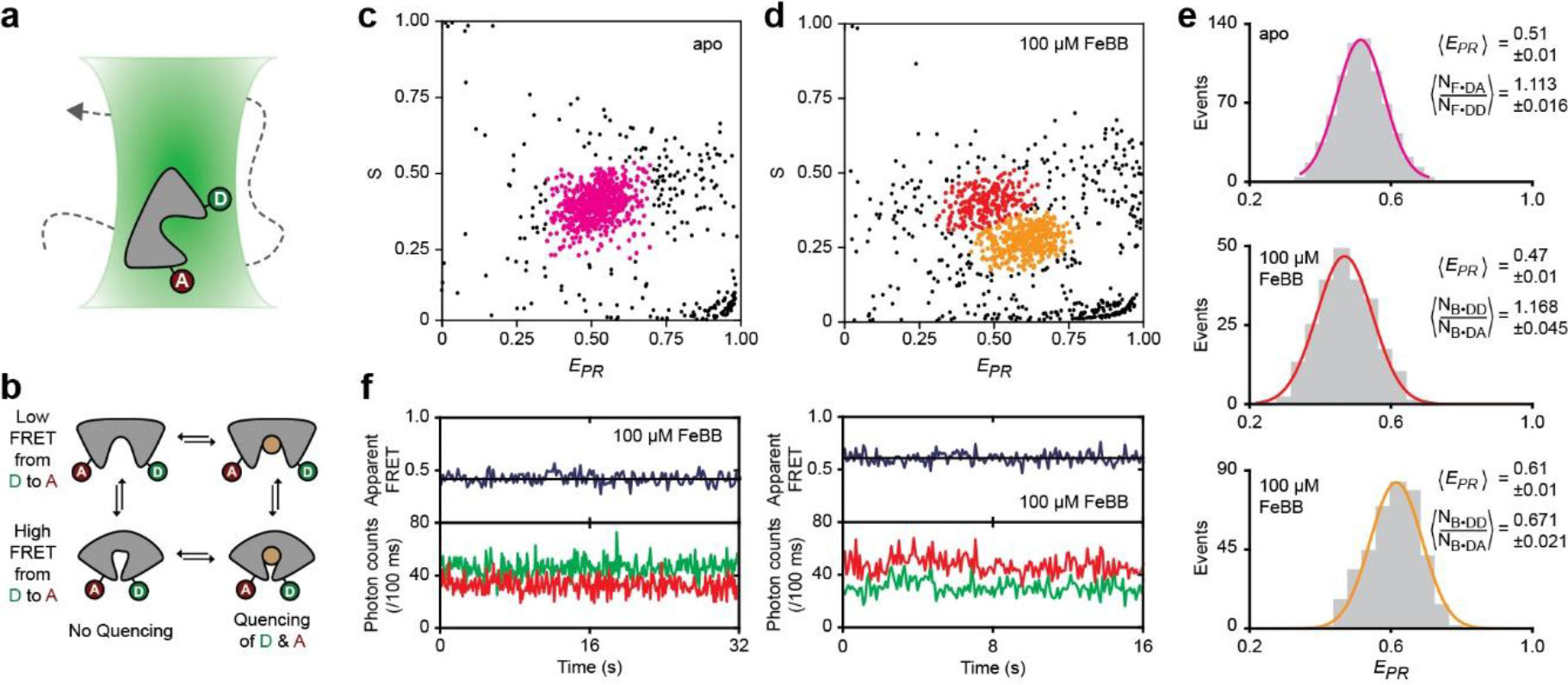
Conformational changes of FeuA in solution. **a**, Cartoon view of labeled proteins diffusing trough the excitation volume of the confocal microscope. **b**, The presence of quenching and FRET depends both on the protein conformation and presence of the ligand FeBB. **c-d**, *E*_*PR*_ and *S* of single FeuA(Q112C/I255C) proteins in the absence (c) and in the presence of 100 μM FeBB (d). **e**,*E*_*PR*_ histogram of the indicated regions of Fig. 3c-d. Solid line is a Gaussian fit. A 95% confidence interval for the average *E*_*PR*_ and count rate rations are indicated. **f**, Representative fluorescence trajectories of surface-immobilized FeuA(Q112C/I255C) in the presence of 100 μM FeBB. Top panel shows calculated apparent FRET efficiency (blue) from the donor (green) and acceptor (red) photon counts as shown in the bottom panels. Black line indicates the average efficiency. The number of analysed molecules is provided in Table S3

The *E*_*PR*_ and *S* values of many individual proteins were acquired in the absence or presence of saturating concentrations of FeBB (100 μM) (Fig. 3c-d). By separating donor-acceptor labelled proteins from the donor- and acceptor-only labelled proteins based on the *S* range, a *E*_*PR*_ histogram was constructed (Fig. 3e). The *E*_*PR*_ histogram of ligand-free FeuA is unimodal and well fitted by a single Gaussian distribution. In the presence of 100 μM FeBB, two populations of donor-acceptor labelled proteins are observed and are centered around different *E*_*PR*_ and *S* values (Fig. 3d-e). FRET analysis of surface-tethered proteins in the presence of 100 μM FeBB, reveals that FeuA does not switch between these FRET states, i.e., fluorescence trajectories are obtained in either FRET state, with no switching between them (Fig. 3f). The cysteine positions in the crystal structure have distinct distances to the ligand binding site. Therefore, the two FRET states most likely arise due to the different donor and acceptor labelling orientations. This is expected from stochastic labelling of two different cysteine positions whichcauses differences in fluorophore quenching by FeBB and thus differences in *E*_*PR*_ and *S* values. Indeed, analysis of individual acceptor- and donor-only labelled proteins shows that the quenching is position and fluorophore dependent (Fig. 4).

**Figure 4.**
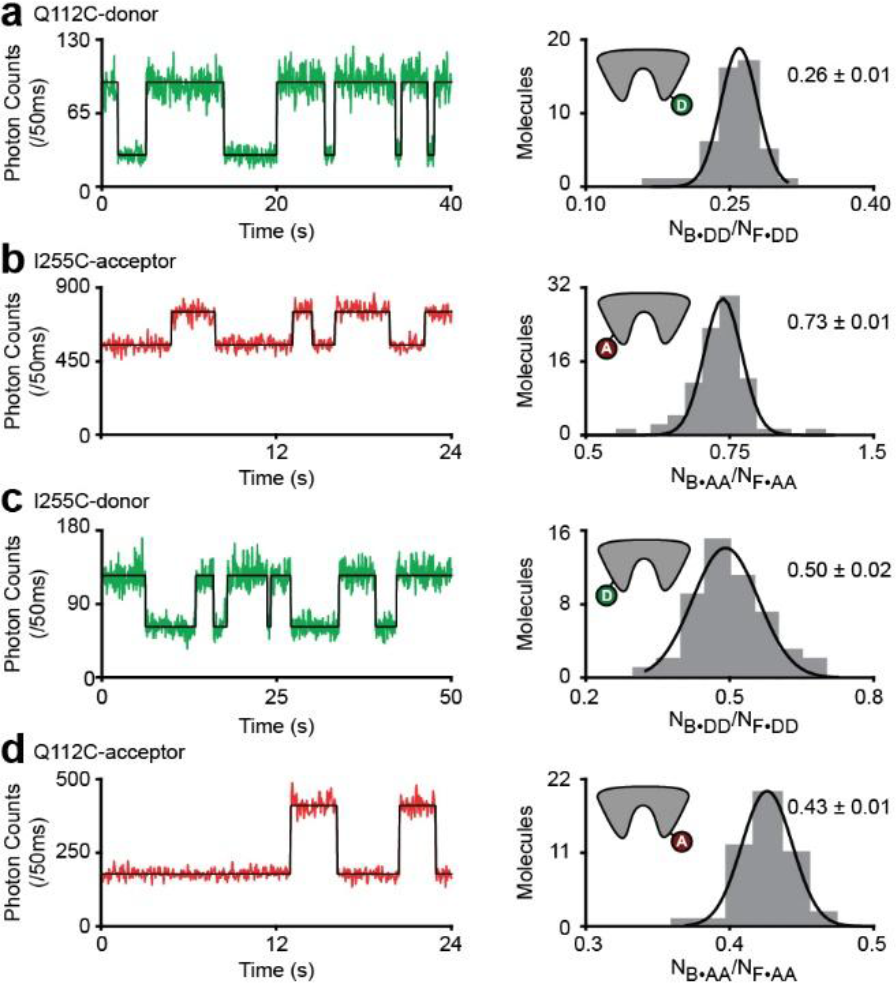
Fluorescence quenching in FeuA depends on the labeling position and the fluorophore. **a-d**, Representative fluorescence trajectories (left) of single-fluorophore labeled FeuA in the presence of 25 nM FeBB (a and c) or 40 nM (b and d) and corresponding histogram of the count rate ratios *N*_*B·DB*_/*N*_*F·DD*_ or *N*_*B·AA*_/*N*_*F·AA*_ of all molecules (right). In the fluorescence trajectories: donor (green) and acceptor (red) count rate with the most probable state-trajectory of the Hidden Markov Model(HMM) (black) are shown. In the histogram: bars are the data and solid line a Gaussian fit. A 95% confidence interval for the average of the brightness ratios are indicated. The number of analysed molecules is provided in Table S3.

To correct the FRET efficiencies for fluorophore quenching, we related the populations in Fig. 3d to its corresponding labelling orientation. The low *S* value of the high FRET population in Fig. 3d (orange population) implies that the quenching of the donor is more prominent than that of the acceptor. To quantify the quenching we prepared and studied all four single cysteine mutants also used in our FRET assays (Fig. 4). The quenching behavior seen in Fig. 3b was observed when Q112C was labelled with a donor and I255C with an acceptor (Fig. 4a-b). The largely unaltered *S* value of the low FRET population in Fig. 3d (red population) suggests that donor and acceptor quenching is similar and was observed to occur when the labelling orientation is reversed, i.e., Q112C is labelled with an acceptor and I255C with a donor (Fig. 4c-d).

To evaluate whether ligand binding conformational changes in FeuA are coupled, we developed an analysis scheme that describes the influence of (i) donor and acceptor quenching (by ligand FeBB) and (ii) considers FRET between the donor and acceptor fluorophores. We show in the Materials and Methods section and Supplementary Information that

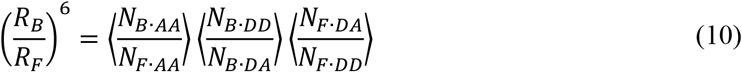

is an unbiased and consistent estimator for (*r*_*B*_/*r*_*F*_)^6^, where *r*_*B*_ and *r*_*F*_ are the interprobe distance of the ligand-bound (B) and ligand-free (F) proteins. In equation (10), *N*_*i·XY*_ denotes the measured count rate of *Y* emission (<u>D</u>onor, <u>A</u>cceptor) upon *X* excitation (<u>D</u>onor, <u>A</u>cceptor) when being in state *i* (<u>B</u>ound, <u>F</u>ree) and 〈·〉 denotes the average. Noteworthy, the distance ratio is independent of donor quenching, i.e. of *N*_*B·DD*_/*N*_*F·DD*_.

The average ratios *N*_*B·DD*_/*N*_*F·DD*_ and *N*_*F·DA*_/*N*_*F·DD*_ are obtained from the selected molecules in Fig. 3c-d and noting that *E*_*PR*_ = *N*_*i·DA*_/(*N*_*i·DA*_ + *N*_*i·DD*_) (Fig. 3e). The average ratio *N*_*B·AA*_/*N*_*F·AA*_ is obtained for individual acceptor-only labelled proteins and by determining the ratio between the acceptor fluorescence intensity in the free and ligand-bound state (Fig. 4b, d). All values used for the distance ratio estimation are provided in Table 1. Finally, by using equation (10), we find that the distance ratio *r*_*B*_/*r*_*F*_, when Q112C is labelled with donor and I255C with acceptor, is estimated to be 0.90 ± 0.01 (95% confidence interval (CI)) and remains the same (0.91 ± 0.01) when the labeling orientation is revered (Table 1). These values are in good agreement with those calculated from the crystal structures of ligand-free and FeBB-bound FeuA^20^ that predict a 86% reduction in C_α_-C_α_ distance between the residues Q112 and I255. In summary, the analysis reveals that under FeBB induces a conformational change in FeuA in buffer solution. Moreover, the reduced interprobe distance in the presence of ligand is consistent with the view that the conformational transition from the open to the closed state in FeuA and related SBPs^4^ are driven by ligand-protein interactions. However, the precise ligand-binding mechanism and whether there are short-lived intermediate states or fast conformational sampling cannot be concluded from these measurements.

**Table 1.** Interprobe distance ratio estimation.

| Assumed labelling<br>orientation | $\left\langle \frac{N_{B\cdot AA}}{N_{F\cdot AA}} \right\rangle$ | $\left\langle \frac{N_{B\cdot DD}}{N_{B\cdot DA}} \right\rangle$ | $\left\langle \frac{N_{F\cdot DA}}{N_{F\cdot DD}} \right\rangle$ | $\left( \frac{R_B}{R_F} \right)^6$ | $\frac{R_B}{R_F}$ |
| --- | --- | --- | --- | --- | --- |
| Q112C-I255C<br>donor-acceptor | 0.733 ±<br>0.010 | 0.671 ±<br>0.021 | 1.113 ±<br>0.016 | 0.547 ±<br>0.020 | 0.904 ±<br>0.006 |
| Q112C-I255C<br>acceptor-donor | 0.432 ±<br>0.008 | 1.168 ±<br>0.045 | 1.113 ±<br>0.016 | 0.562 ±<br>0.025 | 0.908 ±<br>0.007 |

Error denotes a 95% confidence interval.

### Rare intrinsic conformational transitions

To directly observe how binding and unbinding of the ligand FeBB are coupled to conformational changes, and to obtain insight into conformational dynamics, individual proteins were studied for extended times by investigating surface-tethered FeuA proteins with smFRET. It is worthwhile to note that in our surface-based smFRET assays we do not determine absolute distances, but only monitor relative distance changes by determining the instrument-dependent apparent FRET efficiency.

With this, we investigated the dynamics of ligand-free FeuA and addressed whether the protein can also close intrinsically, i.e. when the ligand is absent. Compared to the solution-based smFRET experiments, examining individual surface-tethered proteins greatly increases the sensitivity to detect rare events. To investigate a truly ligand-free protein a high concentration of unlabeled FeuA protein (~20 μM) was added to scavenge any potential ligand contamination that could otherwise cause ligand-induced closing events.

Consistent with the solution-based smFRET measurements (Fig. 3c), FeuA is in a single FRET state (the open conformation) in the majority of fluorescence-trajectories (280 traces out of 294). In these traces, FeuA shows no detectable changes in the apparent FRET efficiency (or fluorescence intensity) related to conformational changes. However, in a small number of trances (14 traces out of 294), we observed rare transitions to a high FRET state, suggesting that FeuA can intrinsically close (Fig. 5). Indeed, by using equation (9) (see Materials and Methods) the average distance ratio between this high and low FRET state is 0.88 ± 0.02 (95% CI). This shows that the interprobe distance of this rare ligand-free protein state is reduced and inferred to be closure of the protein. Importantly, the absence of additional quenching effects provides direct evidence that the conformational change occurs independently of the ligand FeBB. From the analysis of all the transitions, we find that the ligand-free closed state has an average lifetime of 37 ± 9 ms (mean ± s.e.m.) and is formed on average only once every 38 s (Fig. 5). In summary, ligand-free FeuA is primarily in an open conformation and can extremely rarely close, leading to the formation of a short-lived closed conformation.

**Figure 5.**
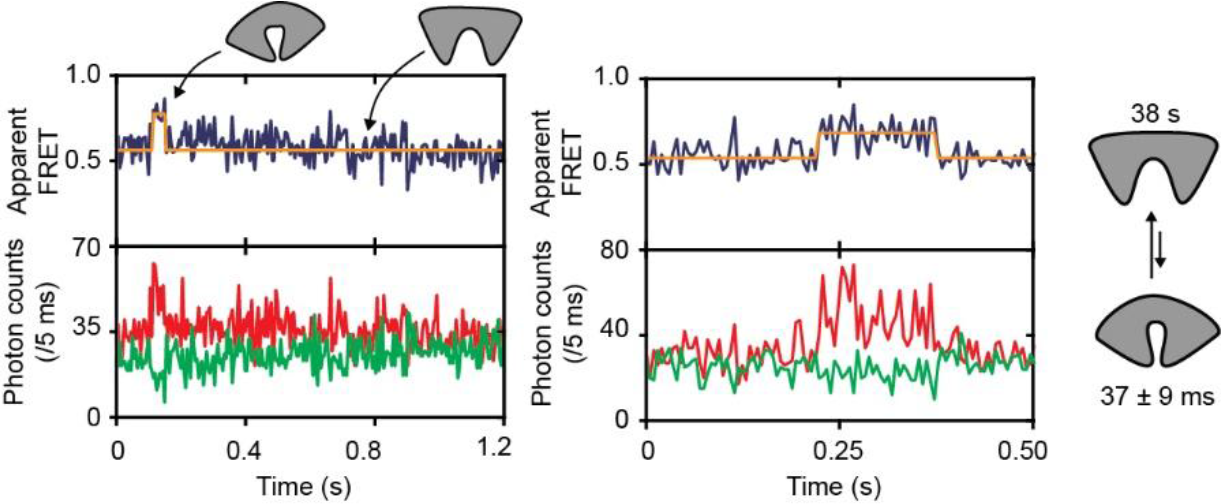
Ligand-free conformational dynamics of FeuA. Representative fluorescence trajectories of surface-immobilized FeuA in the absence of ligand. 20 μM unlabeled FeuA was present to remove potential traces of contaminating ligands. The top panel shows apparent FRET efficiency (blue) which was derived from the donor (green) and acceptor (red) photon count rates shown in the bottom panel. The cartoon depicts the open and closed conformation in the absence of ligands with their respective lifetimes. The number of analysed molecules is provided in Table S3.

### The ligand-bound protein is in the closed conformation

We then investigated the conformational dynamics of the ligand-bound protein. In our assay, the total fluorescence intensity reports on the presence of the ligand, whereas additional apparent FRET efficiency changes are indicative for protein conformational changes. However, in the 105 fluorescence-trajectories that were recorded in the presence of ~K_D_ concentrations of FeBB we could not observe any FRET changes within the period a ligand was bound by FeuA, i.e. the low intensity quenched state (Fig. 6a; Fig. S1a). By examining 140 binding events we observed that the average apparent FRET efficiency was not significantly different within each period as compared to the initial or final 200 ms (*P*=0.28, one-way analysis of variance (ANOVA); Fig. 6b). This suggests that once the ligand is bound, FeuA remains closed and other conformational transitions, such as formation of an open liganded-state, do not occur (or only on timescales faster than 200 ms, see below).

**Figure 6.**
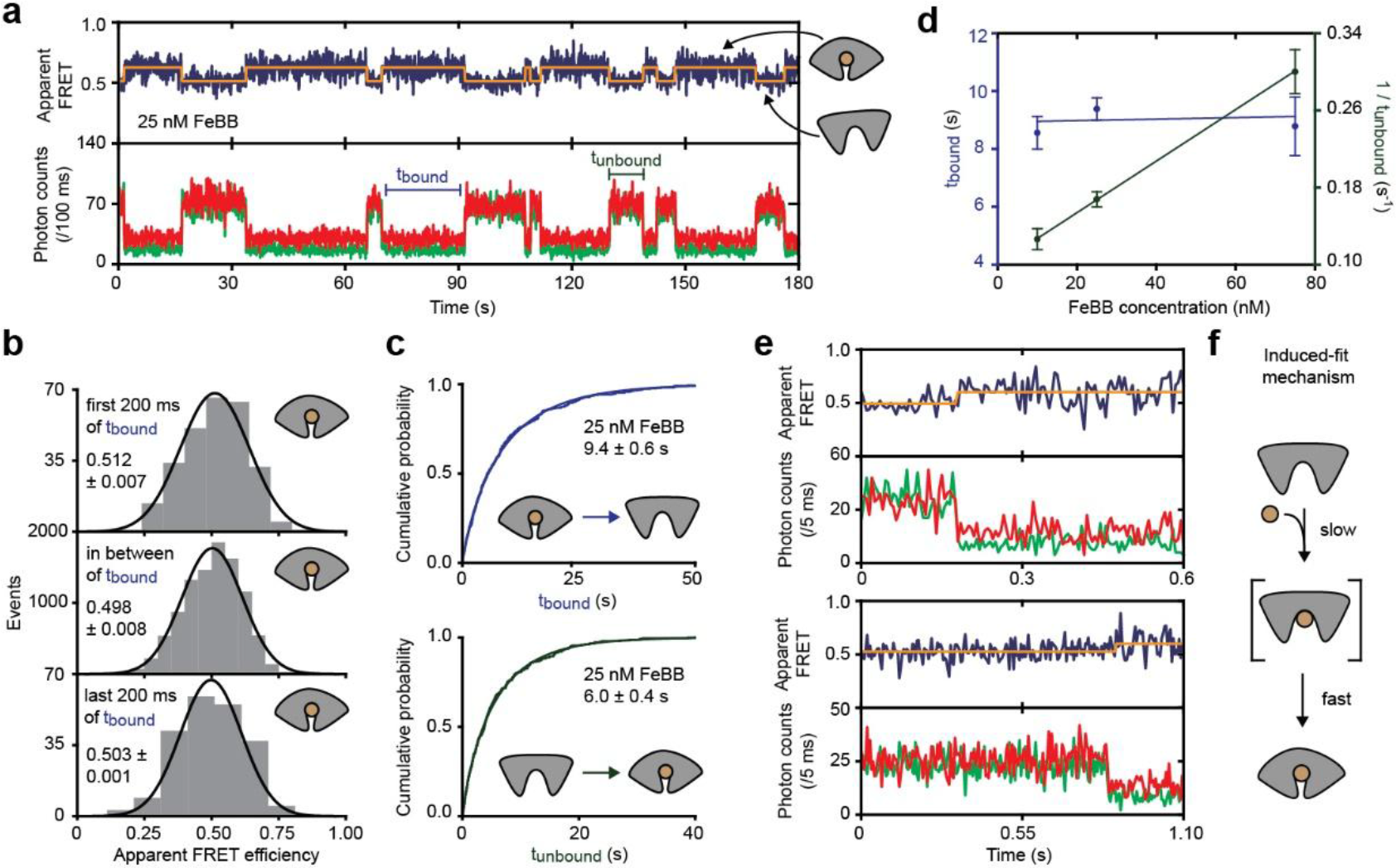
FeuA binds ligand via an induced-fit mechanism. **a**, Representative fluorescence trajectories of FeuA. In all fluorescence trajectories presented in the figure, the top panel shows apparent FRET efficiency (blue) and donor (green) and acceptor (red) photon counts in the bottom panel. Orange lines are the most probable state-trajectory of the Hidden Markov Model (HMM). **b**, Apparent FRET efficiency histogram of the first and last 200 ms of the binding event and the period between that. Mean ± s.e.m. is indicated **c**, Cumulative distribution function (CDF) of the time FeuA has FeBB bound (t_bound_; top) or is free (t_unbound_; bottom). Discontinuous line denotes non-parametric CDF estimate and continuous line an empirical Weibull distribution fit. **d**, t_bound_ (purple) and the rate of binding (1/t_unbound_; green) as function of FeBB concentration. Data is mean ± s.e.m. and solid line a linear fit. From the fit a binding and unbinding rate of 3 10^6^ M^−1^s^1^ and 0.11 s^−1^ are obtained, to yield a K_D_ of 33 nM. **e**, Representative fluorescence trajectories of FeuA showing a binding event. **f**, Schematic of induced-fit mechanism. Number of analysed molecules in Table S3.

With this in mind we determined the lifetime of the ligand-bound state at varying FeBB concentrations. We observed that the lifetime of this state was largely concentration independent (*P*=0.63, one-way ANOVA) and has an average lifetime of 9.0 ± 0.2 s (mean ± s.e.m.; Fig. 6c-d; Fig. S1b). Interestingly, the ligand-bound closed conformation is 250-fold longer lived that the intrinsic closed state (9.0 ± 0.2 s *versus* 37 ± 9 ms).

### Ligand are recognized and bound via an induced-fit mechanism

Next, we investigated which conformational state binds the ligand (intrinsically closed or the open state), and thus whether the ligand is bound via a conformational selection or an induced-fit mechanism (Fig. 1). We expect that when the intrinsic closed conformation binds the ligand, the ligand binding frequency would be limited by the intrinsic closing frequency (~1.6 min^−1^). So we determined the lifetimes of the ligand-free states at varying FeBB concentrations (Fig. 6d; Fig. S1a, c). We observed that the binding/closing frequency increases linearly with FeBB concentration and is already 7.8 ± 0.2 min^−1^ (mean ± s.e.m.) for the lowest concentration measured (10 nM FeBB) (Fig. 6d). Thus ligand binding occurs at a faster rate than the intrinsic closing rate. These data are consistent with ligand-binding occurring via an induced-fit mechanism. In addition, in traces recorded at higher excitation intensity, as a way to increase the time resolution from 100 to 5 ms, shows no substantial FRET changes prior to binding of the ligand, e.g. it reveals the absence of intrinsic closing before the ligand binds (Fig. 6e). The average apparent FRET efficiency of the 10-ms-period before the ligand binds (0.53 ± 0.02, mean ± s.e.m.) and the period prior to that (0.53 ± <0.01, mean ± s.e.m.), when the protein is in the open conformation, are not significantly different (*P*=0.88, two-tailed unpaired *t*-test). This shows that the open conformation binds the ligand and that ligand binding occurs via induced-fit (Fig. 6f).

### Open conformation in complex with ligand is extremely short-lived

An essential intermediate state of the induced-fit mechanism is the open-liganded state (Fig. 1). Based on our data we concluded already that the open-liganded state has to be shorter-lived than 200 ms (see ‘The ligand-bound protein is in the closed conformation’ section). To further investigate the lifetime of this state, we increased the excitation intensity to obtain a time resolution of 4 ms. To probe the open-liganded state, we used saturated amounts of FeBB. Under these conditions the ligand-free open conformation is expected to be absent and any detected open state would consequently correspond to the open-liganded form. However, by examining 94 individual molecules with a total observation time of 104 s, we could not detect any opening transitions (Fig. 7). All these traces show FRET fluctuations, but those could not be separated from noise and did not originate from a clear anti-correlated donor and acceptor fluorescence change, as expected for real changes in FRET efficiency.

**Figure 7.**
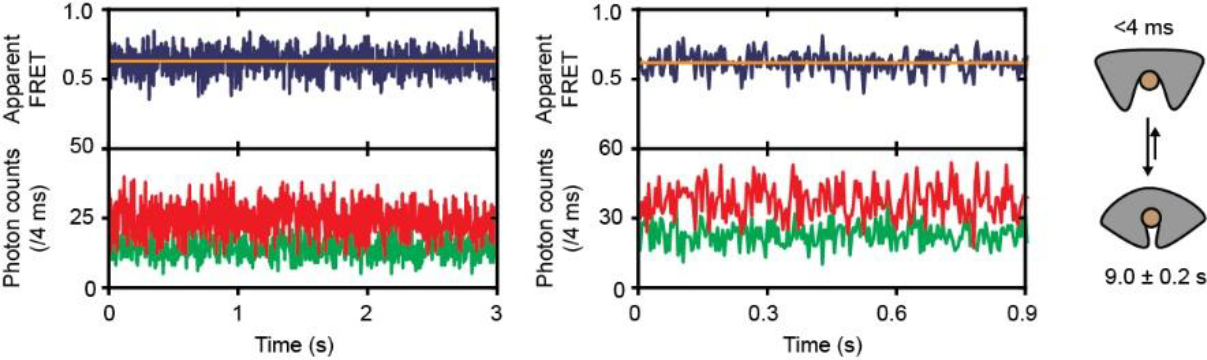
Ligand-bound conformational dynamics of FeuA. Representative fluorescence trajectories of surface-immobilized FeuA(Q112C/I255C) in the presence of 100 μM FeBB. To improve the temporal resolution, the excitation intensity was increased to 5 kW/cm^2^. The top panel shows calculated apparent FRET efficiency (blue) from the donor (green) and acceptor (red) photon counts as presented in bottom panel. Orange line indicate average apparent FRET efficiency value. The cartoon depicts the open and intrinsic closed conformation with its lifetime indicated. The number of analysed molecules is provided in Table S3.

Although we could not directly observe the open-liganded state, we were able obtain an upper bound for its lifetime. Based on the time resolution of the measurement we conclude that the open-liganded state should be shorter lived than 4 ms. Thus, when FeuA is in complex with FeBB, closing is accelerated more than 10000-fold, i.e., from <0.004 s to 38 s. This suggests a mechanism in which the ligand drastically accelerates the conformational change that is already present in the ligand-free protein.

### The energy landscape of FeuA

So far we focused on the dynamical aspect of the molecular recognition process that ultimately originates from the precise architecture of the energy landscape of the combined system comprising the protein and ligand. Here, we use our single-molecule results to determine the thermodynamic properties of FeuA (Fig. 8).

**Figure 8.**
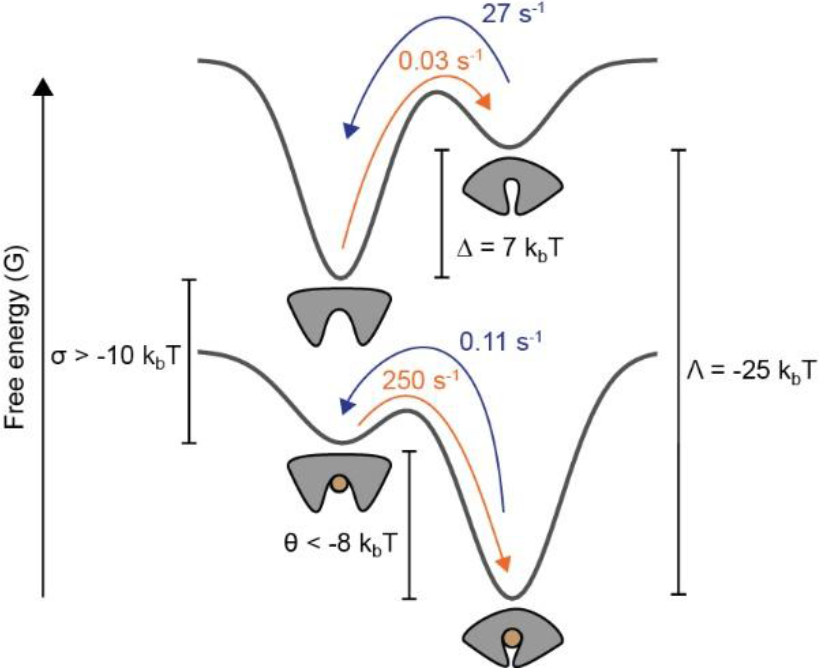
The conformational landscape of FeuA. Schematic representation of the thermodynamics of ligand binding and conformational states of FeuA. Details on the determination of energy values can be found in results section.

The binding process can most easily be treated within the context of Gibbs ensembles. The grand partition function Ω(*T*, *μ*) of a single protein-ligand system as shown in Fig. 8 is

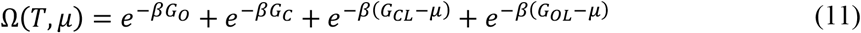

where *β* is (*k*_*b*_*T*)^−1^, *k*_*b*_ is the Boltzmann constant, *T* is the absolute temperature, *G*_*i*_ is the free energy of state *i* (*O* open-unliganded; *C* closed-unliganded; *OL* open-liganded; *CL* closed-liganded) and *μ* is the chemical potential. We assume that the ligand solution can be treated as ideal, so *μ* = *μ*_0_ + *k*_*b*_*T* ln *L*, where *L* is the ligand concentration (relative to 1 Molar) and *μ*_0_ is the standard chemical potential, i.e. *μ* = *μ*_0_ when *L* = 1.

The probability that the protein is in the intrinsic closed conformation is

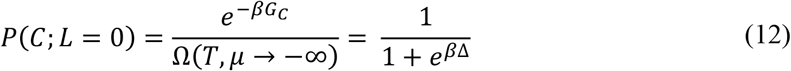

where Δ = *G*_*C*_ − *G*_*O*_ is the ligand-free protein conformational free energy. From the fraction of time spend in the intrinsic closed conformation in the absence of ligand (Fig. 5), we find that *P*(*C*; *L* = 0) is 10^−3^ so Δ = 7 *k*_*b*_*T*.

In the presence of ligand, the fraction of proteins occupying a ligand is given by

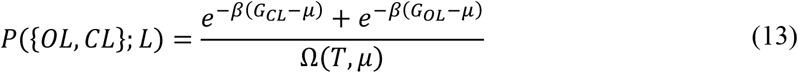

By treating *μ* as an ideal ligand solution and 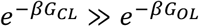 (see also below) we find that equation (13) reduces to

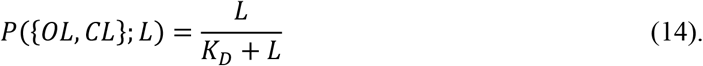

Here *K*_*D*_ is the dissociation constant as determined in our study equal to

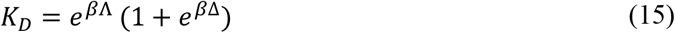

where Λ = (*G*_*CL*_ − *μ*_0_) − *G*_*C*_ is the protein-ligand interaction free energy of the closed conformation. We found that FeuA binds FeBB with a *K*_*D*_ of 20 nM so Λ = −25 *k*_*b*_*T*.

When the protein is saturated with ligand, the probability of being in the closed conformation is

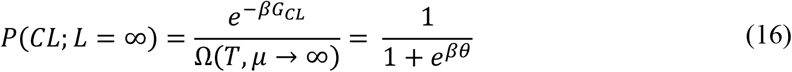

where *θ* = *G*_*CL*_ − *G*_*OL*_ is the ligand-bound protein conformational free energy and *P*(*OL*; *L* = ∞) = 1 − *P*(*CL*; *L* = ∞). We could not directly observe the *OL* state in the smFRET measurements (Fig. 7), but we can calculate an upper bound for *P*(*OL*; *L* = ∞). An estimator for *P*(*OL*; *L* = ∞) is

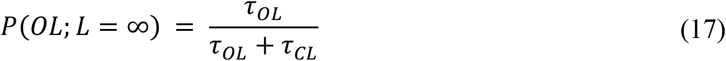

where *τ*_*OL*_ and *τ*_*CL*_ are the lifetime of the *OL* and *CL* state, respectively. By using *τ*_*OL*_ < 4 ms and *τ*_*CL*_ = 9.0 s, we find that *P*(*OL*; *L* = ∞) < 5 10^−4^ so that *θ* < −8 *k*_*b*_*T*. Finally, we find that σ = (*G*_*OL*_ − *μ*_0_) − *G*_*O*_ = Δ + Λ − *θ* > −10 *k*_*b*_*T*, where *σ* is the protein-ligand interaction free energy of the open conformation.

Taken together, in the absence of ligand the closed conformation is thermodynamically unstable and requires an energy input of Δ = 7 *k*_*b*_*T* for its formation. The situation is completely reversed when FeuA has a ligand bound; the open conformation is thermodynamically unstable and requires an energy input of more than −*θ* = 8*k*_*b*_*T* for its formation. From the fact that Λ − σ = (*G*_*CL*_ − *G*_*C*_) − (*G*_*OL*_ − *G*_*O*_) < −15 *k*_*b*_*T* we conclude that the protein-ligand interactions are at least 15 *k*_*b*_*T* stronger in the closed conformation compared to the open conformation.

## DISCUSSION

Modulation of the protein conformational landscape by ligand interactions is fundamental to the function and regulation of proteins. Here, we used a single-molecule approach to investigate the ligand binding process and to understand how this is coupled to the conformational dynamics of the protein. For this, we analyzed the protein conformation and changes thereof via FRET and visualized the presence or absence of a ligand molecule via fluorophore quenching.

From our analysis of FeuA, combined with recent work of others^4, 7, 9, 10, 12^, a picture emerges where ligand-binding does not induce new protein conformations, rather its interaction with ligands modulates the conformational equilibrium (Fig. 8). It appears that ligand binding only alters the free-energies of the equilibrium states and the barriers between them. For FeuA, the equilibrium shifts towards the open conformation in the absence of ligand, with the closed conformation being 7 *k*_*b*_*T* higher in free energy. On the other hand, when FeuA is bound to a ligand molecule, the closed conformation is more than 8 *k*_*b*_*T* lower in free energy than the open conformation. Because of these large free energy differences, it is clear that in FeuA there is a strong, but not an absolute, coupling between ligand binding and protein conformational changes. This shows that nature has fine-tuned the conformational landscape to approximate the behavior of FeuA by a simple binary-switch that is regulated by ligand interactions.

Traditionally the ligand binding process of (monomeric) proteins can be described by different mechanisms, such as the lock-and-key, induced-fit or a conformational selection mechanism. In the classical lock-and-key mechanism no conformational change occur, while in the induced-fit mechanism, ligand interactions induce a conformational change. In the conformational selection mechanism the intrinsic closed conformation would bind the ligand and shift the equilibrium towards closed state. Here, we demonstrate that ligand recognition occurs via an induced-fit mechanism by showing that the open conformation, rather than the intrinsically closed conformation, binds the ligand (Fig. 6). We argue that the conformational landscape provides the required directionality for the induced-fit mechanism. If ligand-binding was to use a conformational selection mechanism, a substantial amount of thermal energy (Δ = 7 *k*_*b*_*T*) would be required to form the ligand-competent, intrinsic closed conformation, rendering the process highly inefficient (Fig. 8). The induced-fit mechanism would be more efficient, as no thermodynamically unfavorable intermediate states need to be formed during the binding process (σ > −10 *k*_*b*_*T*, *θ* < −8 *k*_*b*_*T*; Fig. 8).

By using sensitive single-molecule methods, we observed that ligand-free FeuA can sample a temporally- and thermodynamically-unstable state that is different from the open conformation (Fig. 5). We provide direct evidence that the conformational change occurs independently of the ligand FeBB, as the FRET change occurs without any additional quenching effects that occurs when the ligand binds. However, further investigations would be required to characterize this conformation on a molecular level, but based on the reduced interprobe distance relative to the open state we infer that this rare protein state represents a closed conformation. We also note that we cannot exclude the existence of other conformations that exchange with the open conformation on the nanosecond-to-microsecond timescale. Such conformations could be other short-lived conformations such as a semi-closed state, as has been observed for the SBP maltose-binding protein^8^; the apparent open state would then be a temporal average of two states. For such scenarios, the binding mechanism would be more complex and might involve ligand binding to a short-lived conformation instead of the open conformation. To further elucidate this, methods with high(er) temporal resolution such as NMR^8^, pulsed interleaved excitation (PIE) spectroscopy^34^ or multiparameter fluorescence detection (MFD)^35^ would be required.

To date, intrinsic closing has only been reported for SBPs of Type I ABC importers^4^. ABC importers that employ SBPs can be subdivided as Type I or Type II based on structural and mechanistic distinctions^36^. FeuA belongs to the Type II FeuBC import system and based on our data, we conclude that intrinsic closing can occur in SBPs of both Type I and Type II systems. Thus, some SBPs^4, 28, 37, 38^, including FeuA, can close both spontaneously and with ligand. In FeuA, ligand interactions drastically accelerate the closing transition more than 10000-fold compared to the intrinsic closing rate (<4 ms *versus* 38 s; Fig. 5; Fig. 7). We speculate that once the open-liganded state is formed, direct ligand interactions pull the domains together, resulting in an acceleration of the closing transition compared to when the ligand is absent.

The ligand does not only accelerate closing, it also temporally stabilizes the conformation by a factor of 250 (Fig. 5; Fig. 6d). Some insight into this temporal stabilization can be obtained from the crystal structures of FeuA^20^. In the holo crystal structure of FeuA the ligand is engulfed by the protein by favorable interactions with the binding-site. Hence it requires substantial input of (thermal) energy to break these interactions and cross the energetic barrier to open the protein again. Contrary, in the intrinsic closed conformation these interactions are not present, allowing a more rapid crossing of the energy barrier.

Taken together, ligand interactions are not necessary for a conformational change in FeuA, however, these interactions accelerate the conformational change (10000-fold) and temporally stabilize the formed conformation (250-fold). Both effects shift in the conformational equilibrium towards the closed conformation. This shift in the conformational equilibrium may have been driven by mechanistic determinants to couple ligand-induced conformational changes in FeuA with transport in FeuBC. Ligand binding by FeuA and related SBPs via an induced-fit mechanism would allow the ABC transporter to discriminate between the ligand-free and ligand-bound states^39^. The ligand-bound FeuA can be used to sense the presence of the correct ligand and initiate the transport process. Some SBPs have additional roles, as they are known to interact with chemoreceptors^40^. Switching between two conformations allow these proteins to transduce a signal, which is allosterically regulated by the ligand. Furthermore, we speculate that a wasteful conversion of chemical energy is prevented by the transient nature and high free energy of the intrinsic closed conformation, as any thermally driven mimic of the ligand-bound FeuA complex might be able to initiate the translocation cycle and consume the energy of ATP. In addition, a competition between the ligand-free and ligand-bound closed conformations to interact with the membrane-embedded transporter would inhibit substrate import into the cell^28^.

As a final comment, we note that our data analysis approach to derive the distance ratio of two (conformational) states with altered quantum yield of donor/acceptor dye could also be applied for situations where FRET is changed due to protein-induced fluorescence enhancement (PIFE)^41–43^. The approach suggested here is particularly attractive for PIFE, since the distance ratio is independent of the donor quantum yield, and thus Cy3, which is the most popular dye for PIFE, could be used in a straightforward fashion without additional knowledge of its quantum yield (changes).

## CONCLUSION

We designed a single-molecule assay and data analysis procedure to probe the FeuA conformational changes via FRET and the presence of the ligand FeBB via fluorophore quenching. We show that FeuA exists in an open and a closed conformation in solution. In the absence of ligand FeuA is (predominately) in the open state and ligand shifts the equilibrium towards the closed state. Ligand binding occurs via the induced-fit mechanism, that is, ligands bind to the open state and subsequently triggers a rapid closing of FeuA in less than 4 ms. Unbinding of the ligand also occurs almost simultaneously with the opening of the protein. However, FeuA also rarely samples the closed conformation without the involvement of the ligand and shows that ligand interactions are not required to close. However, such interactions accelerate the closing transition 10000-fold and decrease the openings rate 250-fold.

## ACKNOWLEGDEMENTS

This work was financed by an NWO Veni grant (722.012.012 to G.G.) and an ERC Starting Grant (No. 638536 – SM-IMPORT to T.C.). G.G. also acknowledges an EMBO fellowship (long-term fellowship ALF 47-2012 to G.G.) and financial support by the Zernike Institute for Advanced Materials. G.G. is a Rega foundation post-Doctoral fellow. Y.A.M. was supported by the Indonesia Endowment Fund for Education (LPDP PhD fellowship to Y.A.M.). T.C. was further supported by the Center of Nanoscience Munich (CeNS), Deutsche Forschungsgemeinschaft within GRK2062/1 (project C03/1) and SFB863 (project A13), LMUexcellent and the Center for integrated protein science Munich (CiPSM).

## AUTHOR CONTRIBUTIONS

M.d.B., G.G., T.C. conceived and designed the study. T.C. supervised the project. G.G. and Y.A.M. performed molecular biology work. M.d.B. performed measurements, analyzed data and developed the theoretical work. M.d.B. prepared the initial draft of the manuscript. M.d.B. and T.C. finalized the manuscript with input of all authors. All authors contributed to the discussion and interpretation of the results.

## AUTHOR INFORMATION

The authors declare no competing financial interest. Correspondence and requests for material should be addressed to T.C.

## SUPPLEMENTARY INFORMATION

### Supplementary Text

The donor and acceptor fluorophore distance ratio of two states, denoted by *r*_1_ and *r*_2_, satisfies (see Materials and Methods section for the full derivation):

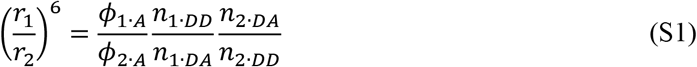

where *n*_*i·DA*_ and *n*_*i·DD*_ are the (background- and spectral crosstalk-corrected) donor and acceptor count rates when the donor is excited and belong to *r*_*i*_. *ϕ*_*i·A*_ is the acceptor quantum yield. Equation (S1) holds when the refractive index of the medium, the dipole orientation factor *κ*^2^, the molar extinction coefficient of the acceptor and the normalized donor emission spectra are the same for state 1 and 2.

Here, we will consider how the distance ratio (*r*_1_/*r*_2_)^6^ can be estimated from the data. We use the following notation: *N*_*i·XY*_ represents the measured count rate of *Y* emission (*<u>A</u>*cceptor, *<u>D</u>*onor) upon *X* excitation (*<u>A</u>*cceptor, *<u>D</u>*onor) belonging to *r*_*i*_. *N*_*i·XY*_ is corrected for spectral crosstalk and background. In the derivation below we assume that the relaxation times of the excited states of the fluorophores are short compared to the time between two consecutively detected photons, so that there is no correlation between consecutive photons and the distribution of *N*_*i·XY*_ can be approximated by a Poisson distribution with parameter *n*_*i·XY*_ ^1^. Then,

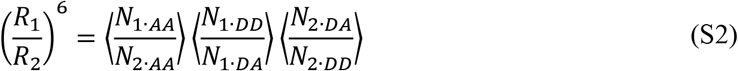

with

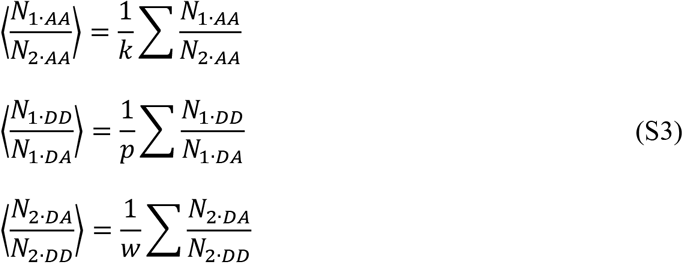

where *k*, *p* and *w* denote the number of observations, is an unbiased and consistent estimator for (*r*_1_/*r*_2_)^6^. The sum in equation (S3) extends over all observations, i.e. the total number of traces or time-bins as described in the main text. Noteworthy, in the absence of additional fluorophore quenching we have *ϕ*_1·*AA*_ = *ϕ*_2·*AA*_ so that

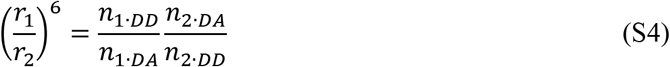

and can be estimated from the data by using the estimator

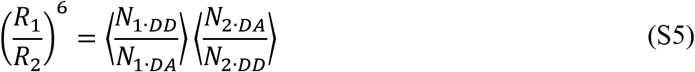

Estimation of the interprobe distance ratio does not require the determination of *γ* or the Föster radius *R*_0_. Below we will focus on the more general scenario as given by equation (S1) and (S2) and note that the results also apply to the more specific case as given by equation (S4) and (S5).

First, we will show that (*R*_1_/*R*_2_)^6^ is an unbiased estimator for (*r*_1_/*r*_2_)^6^ so 𝔼[(*R*_1_/*R*_2_)^6^] = (*r*_1_/*r*_2_)^6^, where 𝔼[*X*] is the expectation value of the random variable *X*. Each term in the product of equation (S2) is independent of each other, so that

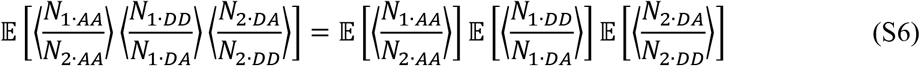

Furthermore, it holds that

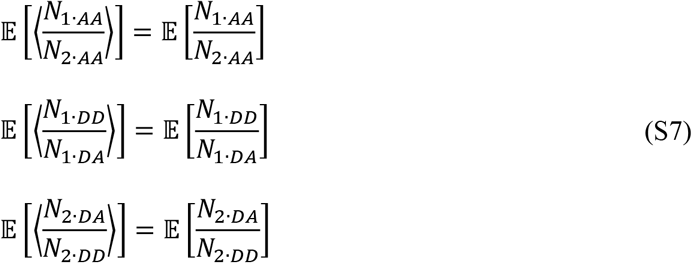

as the terms in the sum of equation (S3) are independent and have the same distribution. By combining equation (S6) and (S7) we have,

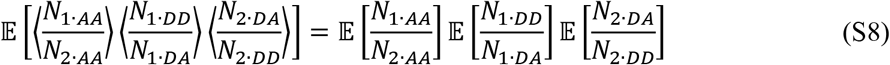

We can approximate each term in equation (S8) further by approximating each term to second-order,

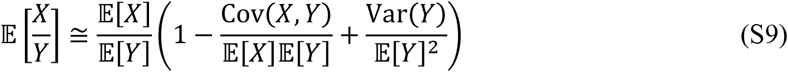

The covariances between *N*_1·*AA*_ and *N*_2·*AA*_, *N*_1·*DD*_ and *N*_1·*DA*_ and of *N*_2·*DA*_ and *N*_2·*DD*_ are zero^2^. Further, under our assumption of Poissonian statistics it holds that Var(*N*_*i·XY*_)/𝔼[*N*_*i·XY*_]^2^ = 𝔼[*N*_*i·XY*_]^−1^ and is thus is negligible when 𝔼[*N*_*i·XY*_] = *n*_*i·XY*_ ≫ 1. Hence, we can safely make the approximation that

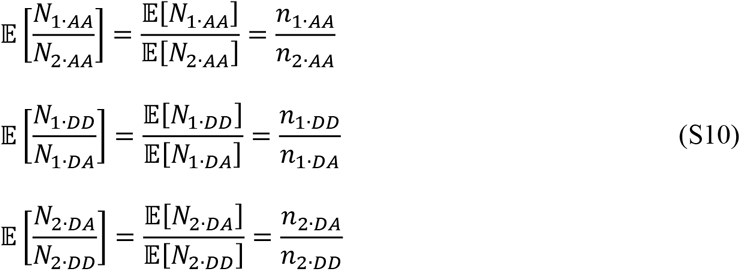

The count rate *n*_*i·AA*_ is the product of the probabilities that (i) the acceptor is excited by the laser (*p*_*EX*_), (ii) the acceptor decays to its ground state by emitting a photon (*ϕ*_*i·A*_) and (iii) the emitted photon is detected (*η*_*A*_)^2^, thus,

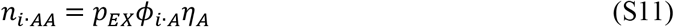

By using equation (S10) and (S11) we have

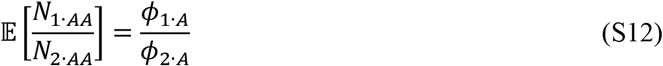

By combing equation (S8), (S10) and (S12) it follows that

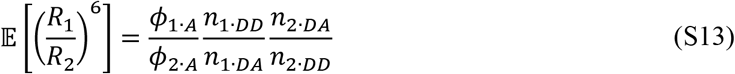

so

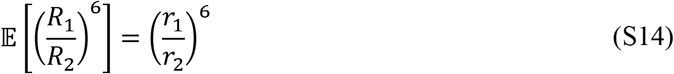

and shows that (*R*_1_/*R*_2_)^6^ is an unbiased estimator for (*r*_1_/*r*_2_)^6^.

If the random variables *X*_*i*_ ··· *X*_*n*_ are independent, then it can be shown that

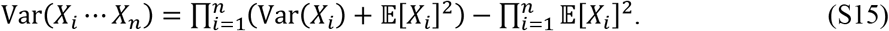

where Var(*X*_*i*_) is the variance of *X*_*i*_. The terms in the product of equation (S2) are independent so by using equation (S15) we find that

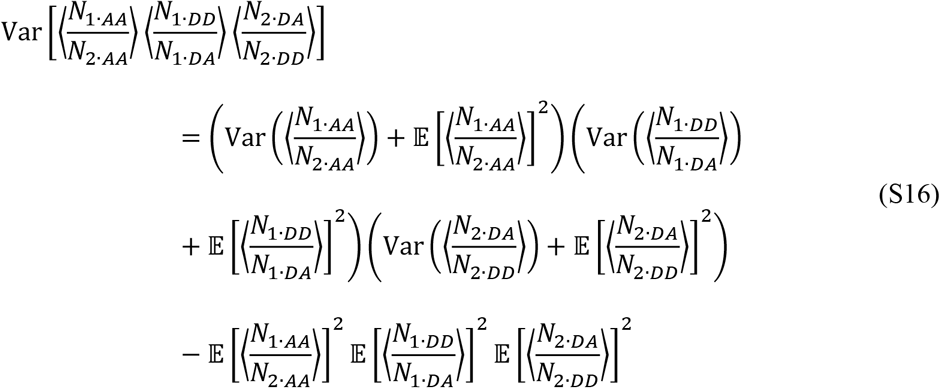

As before, each term in the sum of equation (S3) are also independent and have the same distribution, so

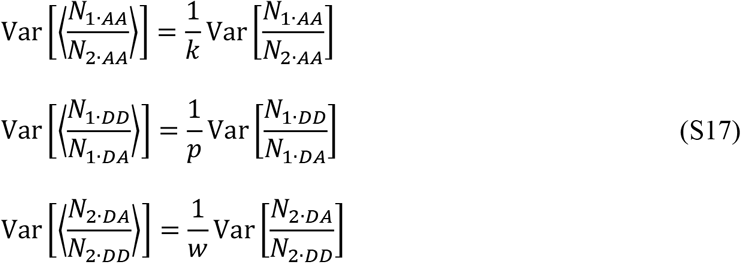

By combining equation (S7), (S16) and (S17) we obtain,

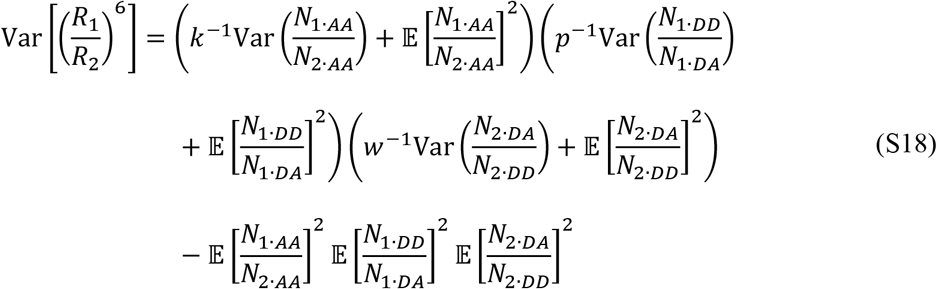

To show that (*R*_1_/*R*_2_)^6^ is a consistent estimator, we need to show that (*R*_1_/*R*_2_)^6^ converges in probability to (*r*_1_/*r*_2_)^6^. We define ***n*** = {*k*,*p*,*l*}, where (*R*_1_/*R*_2_)^6^ depends implicitly on ***n***. We should proof that for any *ɛ* > 0 it holds that,

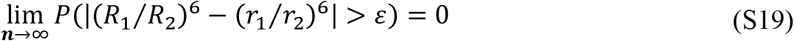

where ***n*** → ∞ should be understood as *k* → ∞, *p* → ∞ and *w* → ∞. By using Chebyshev's inequality and the fact that 𝔼[(*R*_1_/*R*_2_)^6^] = (*r*_1_/*r*_2_)^6^ we can obtain an upper bound for *P*(|(*R*_1_/*R*_2_)^6^ − (*r*_1_/*r*_2_)^6^| > *ɛ*),

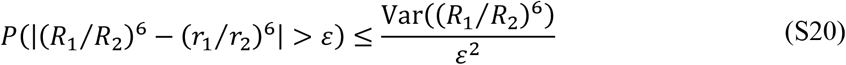

From equation (S18) it follows that

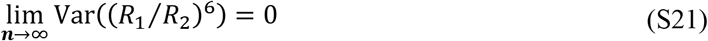

thereby proving that for any ɛ > 0 equation (S19) holds. In conclusion, (*R*_1_/*R*_2_)^6^ is an unbiased and consistent estimator for (*r*_1_/*r*_2_)^6^.

---

**Figure S1.**
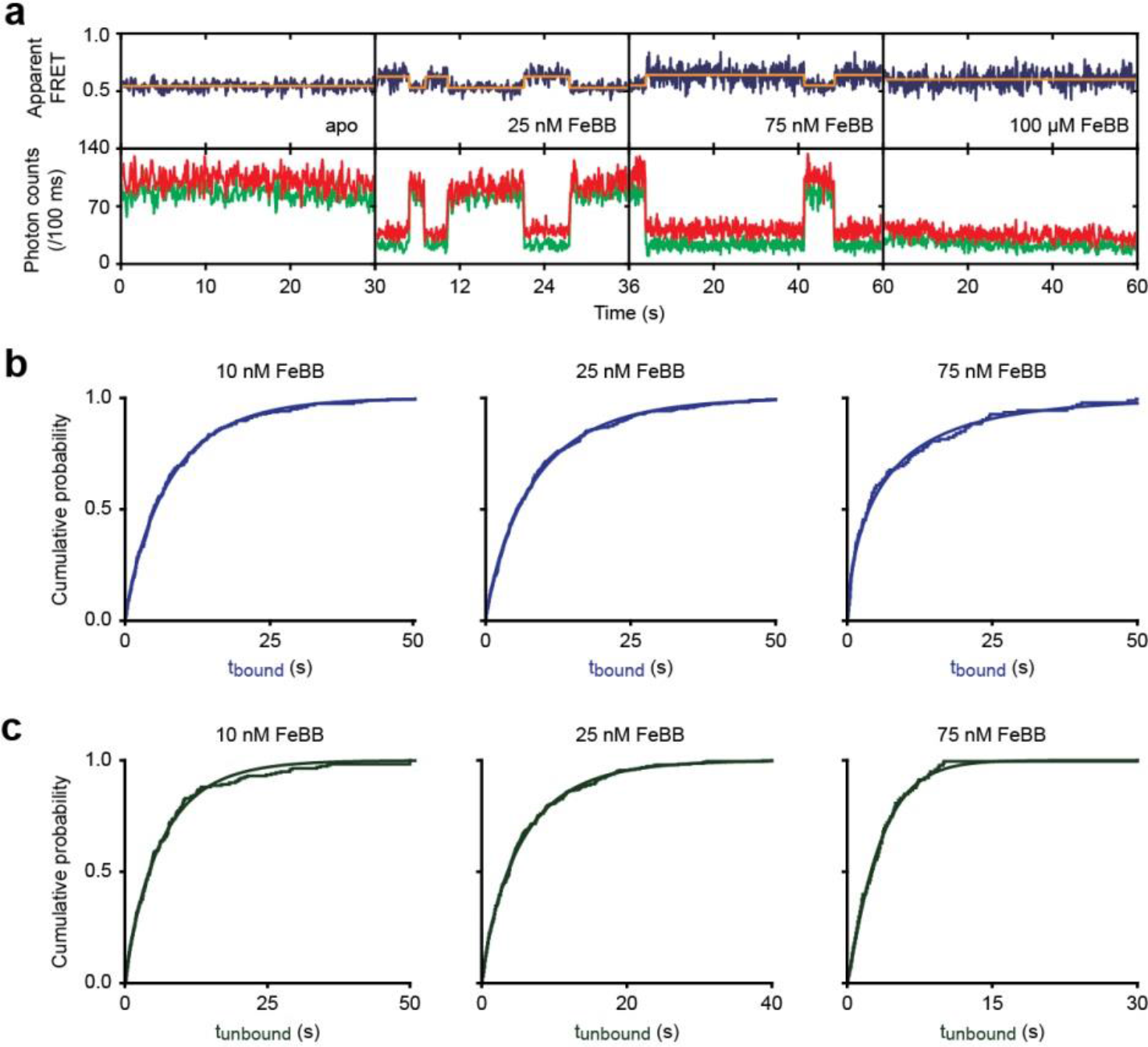
Ligand dependence on protein conformational dynamics and ligand binding kinetics. **a**, Representative fluorescence trajectories of FeuA in the absence and presence of varying concentrations of FeBB as indicated. In all fluorescence trajectories presented: top panel shows calculated apparent FRET efficiency (blue) from the donor (green) and acceptor (red) photon counts as shown in the bottom panels. Orange lines indicates most probable state-trajectory of the Hidden Markov Model (HMM). **b-c**, Cumulative distribution function (CDF) of the time a molecule is bound to FeuA (t_bound_; b) and FeuA is ligand-free (t_unbound_; c). Discontinuous line denotes non-parametric estimate of the CDF and continuous line an empirical fit to a Weibull distribution. Average lifetimes are shown in Fig. 6d.

**Table S1.**
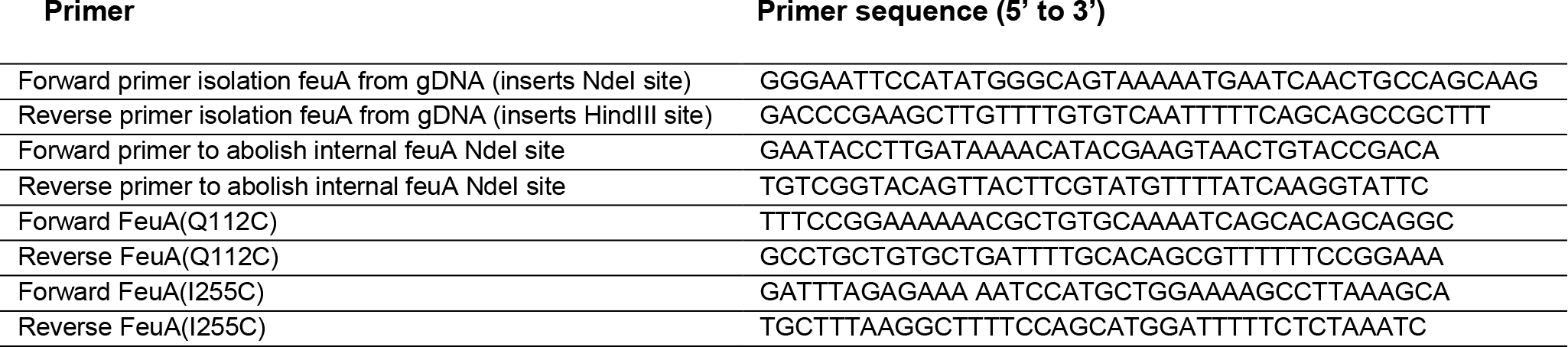
Primers used in this study.

**Table S2.**
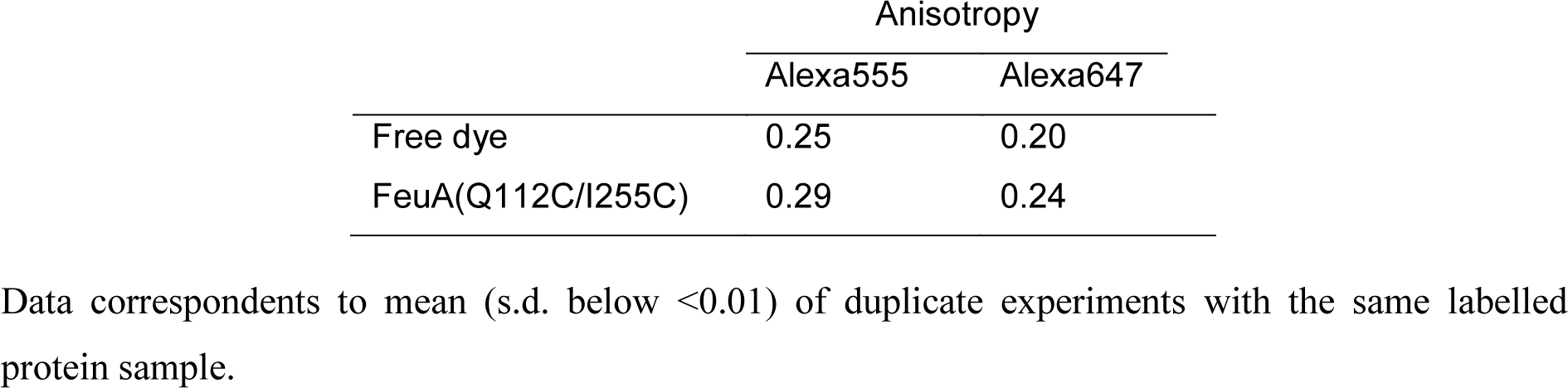
Steady-state anisotropy values.

**Table S3.**
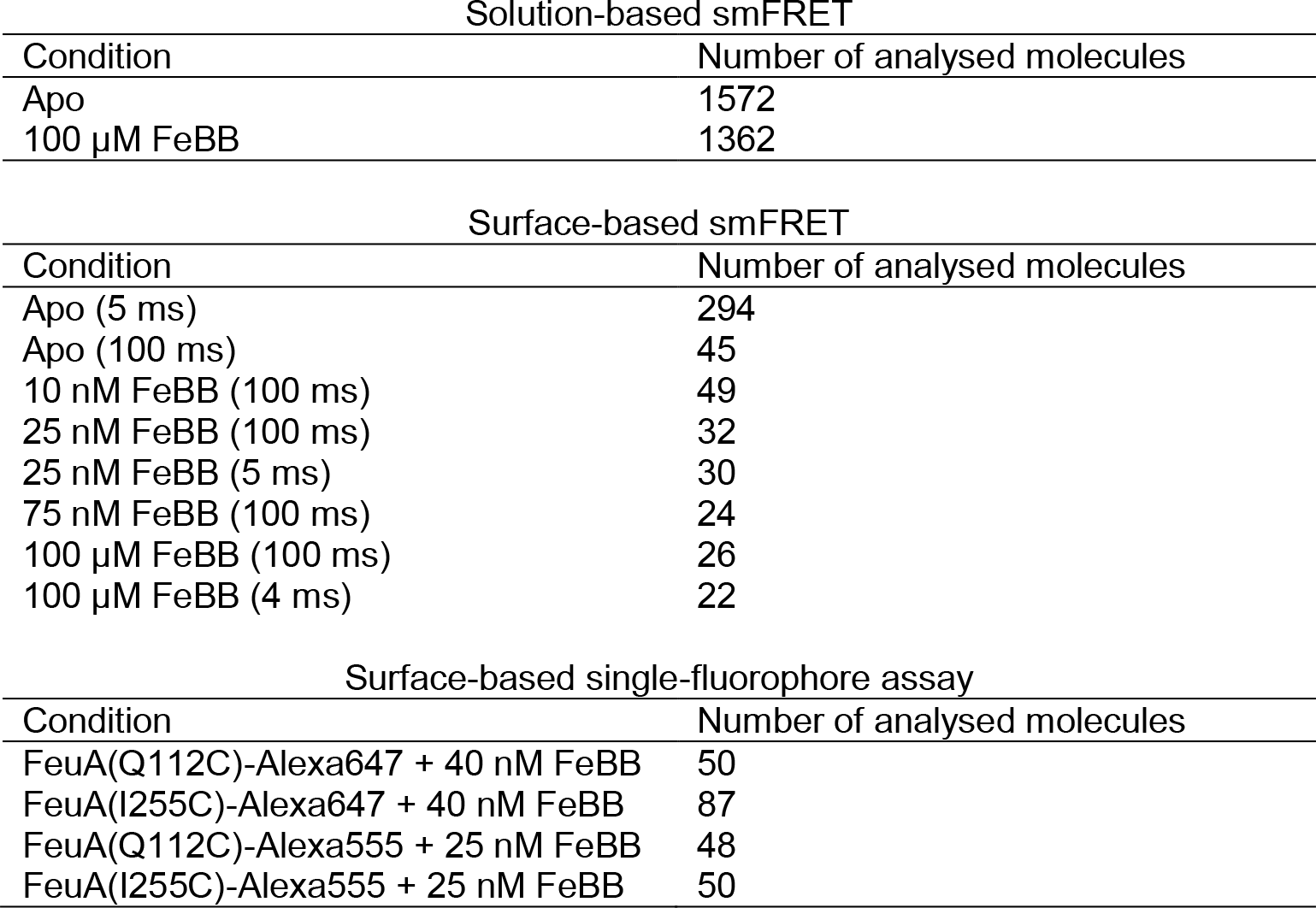
Number of analysed molecules.

| Solution-based smFRET |  |
| --- | --- |
| Condition | Number of analysed molecules |
| Apo | 1572 |
| 100 μM FeBB | 1362 |
| Surface-based smFRET |  |
| Condition | Number of analysed molecules |
| Apo (5 ms) | 294 |
| Apo (100 ms) | 45 |
| 10 nM FeBB (100 ms) | 49 |
| 25 nM FeBB (100 ms) | 32 |
| 25 nM FeBB (5 ms) | 30 |
| 75 nM FeBB (100 ms) | 24 |
| 100 μM FeBB (100 ms) | 26 |
| 100 μM FeBB (4 ms) | 22 |
| Surface-based single-fluorophore assay |  |
| Condition | Number of analysed molecules |
| FeuA(Q112C)-Alexa647 + 40 nM FeBB | 50 |
| FeuA(I255C)-Alexa647 + 40 nM FeBB | 87 |
| FeuA(Q112C)-Alexa555 + 25 nM FeBB | 48 |
| FeuA(I255C)-Alexa555 + 25 nM FeBB | 50 |

